# Mechanistic studies of PFKFB2 reveals a novel inhibitor of its kinase activity

**DOI:** 10.1101/2024.12.25.630325

**Authors:** Craig Eyster, Satoshi Matsuzaki, Atul Pranay, Jennifer R. Giorgione, Anna Faakye, Mostafa Ahmed, Kenneth M. Humphries

## Abstract

The 6-phosphofructo-2-kinase/fructose-2,6-biphosphatase (PFKFB) family of proteins are bifunctional enzymes that are of clinical relevance because of their roles in regulating glycolysis in insulin sensitive tissues and cancer. Here, we sought to express recombinant PFKFB2 and develop a robust protocol to measure its kinase activity. These studies resulted in the unexpected finding that bacterially expressed PFKFB2 is phosphorylated *in situ* on Ser483 but is not a result of autophosphorylation. Recombinant PFKFB2 was used to develop an enzymatic assay to test a library of molecules selected by the Atomwise AtomNet® AI platform. This resulted in the identification of a new inhibitor, B2, that inhibits PFKFB2 (IC_50_ 3.29 μM) and PFKFB3 (IC_50_ 11.89 μM). A-498 cells, which express both PFKFB2 and PFKFB3, were treated with B2. Seahorse XFe analysis revealed B2 inhibited cellular glycolysis and glycolytic capacity. Targeted LC/MS analysis showed B2 decreased fructose-1,6-bisphosphate and downstream glycolytic intermediates but increased fructose-6-phosphate levels, which is consistent with an inhibitory effect on PFK-1 activity. The LC/MS metabolic profile of A-498 cells treated under identical conditions with the known PFKFB3 inhibitor, PFK158, was distinct from that induced by B2. These results thus demonstrate the identification and validation of a new PFKFB kinase inhibitor.

## INTRODUCTION

The 6-phosphofructo-2-kinase/fructose-2,6-biphosphatase (PFKFB) family of proteins are bifunctional enzymes that catalyze the formation or degradation of fructose-2,6-bisphosphate (F-2,6-BP) ^1-3^. F-2,6-BP is an allosteric activator of phosphofructokinase-1 (PFK-1), a committed and rate limiting step of glycolysis. Therefore, depending upon whether F-2,6-BP is being produced or degraded, the dual activities of PFKFB enzymes play a central role in increasing or decreasing glycolysis.

There are four PFKFB isoforms ^2^. In the heart, PFKFB2 is the primary isoform and its kinase activity is enhanced by protein phosphorylation by either Akt, AMPK, or PKA ^1^. Work from our lab has shown that cardiac PFKFB2 is decreased in diabetic mouse models and that its content is regulated by insulin signaling. The constitutive decrease of cardiac PFKFB2 likely contributes to diabetes-induced dysregulation of normal metabolic flexibility cues. Tissue-specific splice variants of PFKFB1 are expressed either in skeletal muscle or liver where they facilitate fine-tuned regulation of glycolysis and/or gluconeogenesis ^1^. In contrast to PFKFB2, protein phosphorylation of PFKFB1 by PKA increases phosphatase activity. PFKFB3 is an inducible member of the PFKFB family and is upregulated in numerous cancers, likely contributing to dysregulation of glucose metabolism and contributing to aerobic glycolysis ^4^. PFKB3 also has a role in other pathophysiological conditions, including liver fibrosis ^5^, sepsis ^6^, endothelial cell response to inflammation ^7^, and physiological processes including adipocyte differentiation ^8^. Lastly, PFKFB4 expression is normally confined to testes but its upregulation is implicated in cancers ^9^.

The central role of PFKFB enzymes in maintaining metabolic flexibility and regulating glucose metabolism make them potential targets for therapeutic intervention. However, progress on this front remains limited in part because of the difficulty in measuring PFKFB enzyme activity, the instability of the kinase product F-2,6-BP, and the lack of available commercial F-2,6-BP standards. With our own interest in the PFKFB2 isoform, we sought to develop a pipeline to measure small molecules that could potentially modulate PFKFB2 and other PFKFB kinase activities. Here we report the recombinant expression, and unexpected serine phosphorylation of PFKFB2 in bacteria, which was subsequently used to optimize an in vitro PFKFB2 kinase activity assay. This assay was then used to screen an AI-screened library ^10^ of small molecules to identify a new inhibitor of PFKFB2 and PFKFB3. We demonstrate that the newly identified inhibitor, which is chemically distinct from known PFKFB inhibitors, is effective in blocking PFK1 activity in vitro. These results thus provide a new PFKFB kinase inhibitor for potentially wide biological applications.

## METHODS

### Recombinant Protein Expression

*PFKFB2 and PFKFB3* constructs (Origene) were subcloned into GST expression vector. Mutation of *PFKFB2* at K174 to G ^1^, H259 to A ^11^, and the double mutant were created using Quickchange (Agilent) methodology. All constructs were transformed into Rosetta cells (Millipore), expressed and induced following standard protocol. Bacteria were harvested via centrifugation and pellets resuspended in 10mL sonication buffer (50mM TrisHCl pH 7.5, 10% glycerol, 150 mM NaCl, 10 mM imidazole, 10 µL 1 M DTT, and 1protease inhibitor tablet (Sigma). Cells were sonicated (power level 5, 45s, 4X, QSonica) at 4°C and spun 10,000 rpm for 10min at 4°C to collect protein lysate. Glutathione-Sepharose beads (Cytiva) were washed twice with PBS and protein lysate was applied and rocked overnight at 4°C. Beads were captured using 10mL column (Biorad) and washed 5X with column volume of PBS. Column was capped and incubated overnight with thrombin in 1mL PBS at 4°C. 1mL PBS was rinsed over the column 3X and protein was concentrated using Amicon Ultra 0.5mL centrifugal filter (3K NMWL) (Millipore). Protein concentration was determined by Nanodrop (Thermo).

### PKA and phosphatase treatments of recombinant protein

Bacterial constructs of PFK2 wild type and mutants were grown overnight in 1ml Terrific Broth/Amp at 37C with shaking. 500ul of overnight culture was added to 10 mL Terrific Broth/Amp. Samples were grown 3 hrs at 37°C with shaking. For induction, 100 µL of 0.04 M IPTG was added to 10 mL TB/Amp. Samples were shaken overnight without heat. Cultures were pelleted and frozen at–80°C. Each pellet was resuspended in 1 mL sonication buffer. 50 mM Tris HCl pH 7.5, 10% glycerol, 150 mM NaCl, 10 mM imidazole, 10 µL 1M DTT, 1 protease inhibitor tablet. A 100 µL sample was taken from each resuspended pellet before sonication to represent 100% phosphorylation. The remaining sample was sonicated in 2x 45s bursts at level 4.5 using a tip sonicator. Assay conditions were 100 µL sonicated sample, 10 µL MnCl_2_ (10 mM NEB), 10 µL phosphatase buffer (10x PMP buffer, NEB) with or without 2 µL Lambda protein phosphatase (NEB, 400000 U/mL). Reactions were carried out for various times. To stop reactions, 30 µL of “stop” buffer mix was added (300 µL Invitrogen SDS 4x sample buffer, 10 µL 1 M DTT, 5 µL Halt Protease and Phosphatase Inhibitor cocktail, Thermo Scientific). All stopped reactions were heated to 95°C for 10 min. Samples were frozen at –80°C for later western blot analysis.

### Western blot analysis

Frozen samples were prepared for SDS PAGE by thawing and sonication (2x 45s). Samples were then diluted 1:4 in 1x sample buffer. 10 µL of each sample was run on SDS page (Invitrogen NP0321) and transferred to a nitrocellulose membrane, blocked for 1 hr with Odyssey TBS blocking buffer (LI-COR). Antibodies used were: PFKFB2 (Cell Signaling, #13029); phosphorylated-PFKFB2 Ser483 (Cell Signaling, #13064); PFKFB3 (Abclonal, A6945); phosphorylated PFKFB3 Ser 461 (ThermoFisher, #PA114619), and anti-GST Tag (Bethyl Laboratories, A190-122A). Primary antibodies were incubated overnight at 4°C, secondary antibody (LI-COR IRDye800CW) was incubated 1 hr at room temperature. Blots were imaged on an Odyssey CLx System and analyzed using the Image Studio Software (LI-COR).

#### AI-Based Small Molecule Virtual Screen

The virtual screen was carried out using the AtomNet^®^ technology, a deep convolutional neural network for structure-based drug design ^12,13^. A single global AtomNet^®^ model was deployed to predict the binding affinity of small molecules to PFKFB2. Details regarding the AtomNet^®^ model training were previously published ^14^. The PFKFB2 crystal structure was used to screen potential inhibitors (PDB ID 5HTK, ^15^). The binding site was specified using the bisphosphatase binding site. This included residues GLU326, TYR337, ARG351, LYS355, TYR366, GLN392, and ARG396.

The Mcule small-molecule library version v20180817, containing ∼10 million small organic molecules for drug discovery purchasable from the chemical vendor Mcule (Palo Alto, CA), was screened. The library in simplified molecular-input line-entry system (SMILES) format was downloaded from Mcule’s website (https://mcule.com/). Every compound in the library was pushed through a standardization process, including removing salts, isotopes, and ions, and conversion to neutral form, conversion of functional groups and aromatic rings to consistent representations. Filters were then applied to some molecular properties, including molecular weight between 100 and 700 Da, the total number of chiral centers in a molecule ≤ 6, the total number of atoms in a molecule ≤ 60, the total number of rotatable bonds ≤ 15, and only molecules containing C, N, S, H, O, P, B, halogens were allowed. Other filters such as toxicophores, Eli Lilly’s MedChem Rules ^16^, and pan-assay interference compounds (PAINS) were also applied to remove compounds with undesirable substructures, resulting in a filtered library of 6,922,894 compounds. A set of 64 poses within the binding site was generated for each small molecule. The trained model scored each pose, and the molecules were ranked by their scores. The top 5000 ranking compounds were examined, and 64 compounds containing diverse chemical scaffolds were selected and obtained from Mcule.

#### PFK-2 kinase activity assay

Relative *in vitro* PFK2 activities were assessed by the levels of fructose-2,6-bisphosphate (F-2,6-BP) produced by either 2.0 µg/mL of purified PFKFB2 (for plate-reader screening) or 5-20 µg/mL PFKFB2/PFKFB3 (for IC50 analyses) incubated in PBS on ambient temperature for one hour in the presence of 5 mM Mg^2+^, 1 mM ATP, and 1mM fructose-6-phosphate (F-6-P). Resultant F-2,6-BP levels were determined by pyrophosphate– fructose-6-phosphate phosphotransferase (PPi-PFK) activity as described previously ^17,18^ with minor modifications. Briefly, after the preincubation phase, the aforementioned F-2,6-BP/PFK2 mixture was diluted 10x by 50 mM pH 8.0 Tris assay buffer, supplemented with 5 mM Mg^2+^, 0.5 mM PPi and 1 mM F-6-P. The PPi-PFK activity was measured as the rate of NADH oxidation (ε_340_ = 6200 M^−1^cm^−1^) following the addition of 150 μM NADH to a mixture of excess PPi-PFK (enriched from potato tubers as described in ^17^), 0.1 μg/mL triosephosphate isomerase (Sigma), 10 μg/mL glycerol-3-phosphate dehydrogenase (Lee BioSolutions), and 0.01 U/mL aldolase (Sigma). The first 10 min of the reaction is an equilibration period and was discarded from rate calculations. PPi-PFK is the rate limiting step of this multi-enzyme reactions.

#### Cell culture and Seahorse XFe analysis

Human kidney carcinoma cell line A-498 (HTB-44) was obtained from ATCC. The effect of PFKFB2/3 inhibitors on glycolysis was determined by glycolysis stress test utilizing Seahorse XFe24 Extracellular Flux Analyzer (Agilent) measuring extracellular acidification rate (ECAR) at basal and maximal glycolysis. After cells reaching confluency in regular DMEM culture media, the media was replaced with either DMSO, 5 µM B2, 25 µM B2, or 5 µM PFK158-supplemented Seahorse media, which is comprised of basal unbuffered XF DMEM, 1 mM sodium pyruvate, and 2 mM glutamine (pH 7.4). During the assay, the following inhibitors were injected sequentially, as is standard for the glycolysis stress test: 10 mM glucose, 1 µM oligomycin, and 50 mM 2-deoxyglucose. After the Seahorse assay, without removing media, 1 µM of Hoechst 33342 was added to each well and subsequently the live cells were counted by Cytation 5 (Aligent) DAPI-channel fluorescent imaging for normalization purpose.

#### Targeted metabolomics of glycolytic intermediates

After A-498 cells reached 95% confluent in 75cm^2^ culture plates, culture media was replaced with DMEM containing either DMSO, 5 µM B2, or 5 µM PFK158. After 60 min incubation at 37°C cells were once washed with warmed PBS and then quickly snap-frozen in LN2. Frozen cell culture plates, stored at – 80°C, were taken out on dry ice/liquid nitrogen and 1.0 mL of chilled methanol (stored at –80°C) was immediately added to the plate and plates were gently swirled for 1 minute on ice to make sure that all the cells were completely covered by methanol. Cells from the plates were then removed with a cell scraper and transferred to a 2.0mL Eppendorf microcentrifuge tube. An additional 200µL of chilled methanol was added to the plate and scraping was repeated for complete recovery of cells. Cells were then sonicated in a water bath for 5 min and then incubated for 20 min on dry ice and transferred to ice for thawing. 350µL milli Q water was then added to the sample and shaken in a thermomixer for 10 min at 4°C. Cells were centrifuged at 15,000 rpm for 10 min and the supernatant was collected. Pellets were stored at –20°C until use for protein estimation. The supernatant was filtered using Agilent Captiva EMR-lipid filter and filtrate was dried in a vacuum centrifuge for approximately 4 hours or until filtrate was completely dried. Dried samples were stored at –80°C and then reconstituted in 100µL of 7:3 v/v acetonitrile and water and transferred to LC/MS vials.

LC-MS analysis was performed on an Agilent 6546 LC/Q-TOF coupled to an Agilent 1290 Infinity II LC. Chromatographic separation was performed on an Agilent InfinityLab Poroshell 120 HILIC-Z, 2.1 × 150 mm, 2.7μm column, coupled with a UHPLC Guard, HILIC-Z, 2.1 mm × 5 mm, 2.7 μm, at 15°C, with a total run time of 29 min. The mobile phase consisted of 20mM ammonium acetate buffer in water (pH=9.2) (A) and acetonitrile (B). To ensure a constant concentration during gradient elution, the InfinityLab deactivator additive (p/n 5191-4506) was added to the aqueous mobile phase. A linear gradient elution was used at a flow rate of 0.4mL/min with the following program: 0.0 min (15% A and 85% B), 1.0 min (15% A and 85% B), 8 min (25% A and 75% B), 12.0 min (40% A and 60% B), 15.0 min (80% A and 20% B), 18.0 min (80% A and 20% B), 19.0 min (15% A and 85% B) and 29.0 min (15% A and 85% B). Data was acquired in a negative ESI full MS scan mode (scan range: m/z 40 to 1,000) using Agilent MassHunter Acquisition software version 10.0. Optimum values for MS parameters were as follows: gas temperature 300°C; drying gas flow 13L/min; Nebulizer pressure 40psi; Sheath gas temperature 350°C; Sheath gas flow 12L/min; Capillary voltage, 3500V; Nozzle voltage 0V; skimmer offset, 45V; Fragmentor 125V; Octopole 1 RF Voltage 750V.

Data analysis and peak integration for target metabolites were performed with Agilent MassHunter Quantitative analysis software (V10.1). An analytical standard mix of glycolytic and TCA cycle intermediates was analyzed along with samples for identification of the metabolites in the samples. For identification of target metabolites in the samples, a custom Agilent Personal Compound Database and Library (PCDL) of target metabolites (glycolytic and tricarboxylic acid cycle intermediates) was created from Agilent METLIN PCDL. The retention times derived from standard mix analysis were also included in the custom PCDL to facilitate the identification of metabolites in the samples. Relative abundance values for target metabolites were obtained by normalizing the raw data with total protein estimated from cell pellets obtained after methanol: water extraction using a Bradford protein assay (Pierce).

#### Statistical Analysis

Statistical analyses were performed on GraphPad Prism Version 10.2.0. IC50 values were derived from least squares fit with [Inhibitor] vs. response (three parameters) using GraphPad Prism version 10.2.0 for Windows, GraphPad Software, Boston, Massachusetts USA, www.graphpad.com. Mass spectrometry peak intensity data was normalized by log transformation (base 10) and mean centering and the default MetaboAnalyst 6.0 settings were used for the principal component analysis and heatmap generation^19^.

## RESULTS

### Bacterially expressed PFKFB2 is phosphorylated *in situ*

Our initial goal was to express PFKFB2 to obtain enzymatically active protein that could be used for subsequent drug screens. PFKFB2 was bacterially expressed as a thrombin-cleavable GST fusion protein (**Supplement Figure 1A**). While most of the PFKFB2 was insoluble, sufficient enzyme was enriched via a GSH-column and thrombin cleavage for subsequent experiments. **Figure 1A** shows total cleared bacterial lysates probed with anti-GST antibody. **Figure 1B** shows the enrichment of recombinant PFKFB2. In eukaryotes, PFKFB2 can be phosphorylated by several different serine/threonine kinases, including AKT, PKA, and AMPK ^1^. This phosphorylation results in enhanced PFKFB2 kinase activity. Western blot analysis revealed that recombinant PFKFB2 purified from bacteria was phosphorylated at Ser483 (**Figure 1A-C**). Phosphorylation was labile, though, with dephosphorylation occurring upon enrichment (**Figure 1B**) and cleavage of PFKFB2 from the GST tag. However, purified PFKFB2 could then be rephosphorylated by treatment with exogenous PKA (**Figures 1B&C**) ^5,20^.

**Figure 1:**
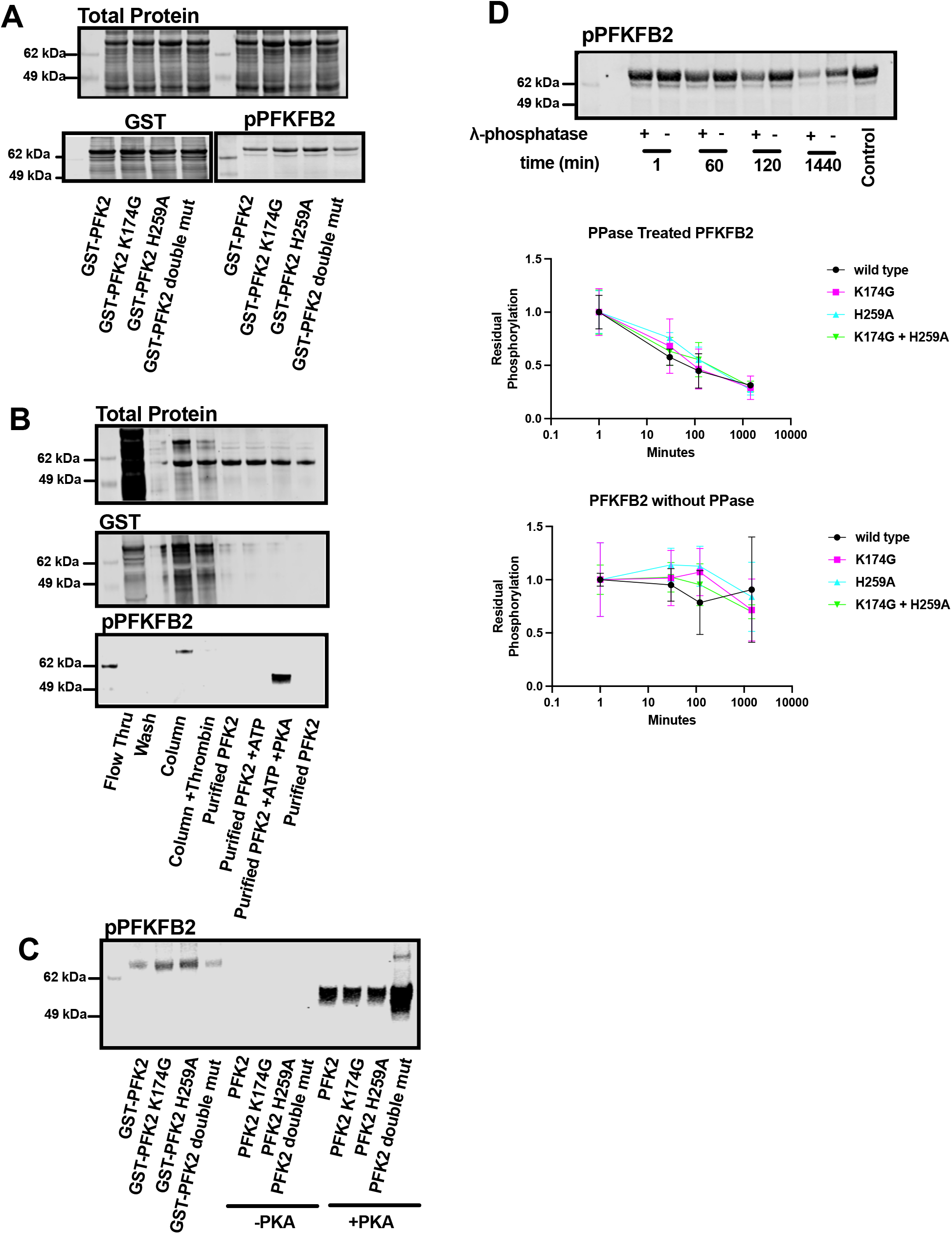
Recombinant PFKFB2 is phosphorylated *in vitro*. (**A**) Western blot analysis of cleared bacterial lysates of GST-PFKFB2 and mutants. Top: Total protein staining of cleared bacterial lysates over-expressing GST-PFKFB2 and mutants. Bottom: Western blots against GST and phosphorylated-PFKFB2 (Ser483) for GST-PFKFB2 and mutants. “Double mut” indicates protein containing both K174G and H259A. (**B**) Western blot analysis of GST-PFKFB2 expression, purification, and *in vitro* phosphorylation. *Flow thru* indicates proteins unbound to GSH beads. *Column* are proteins bound to the GSH beads. *Column + Thrombin* are proteins remaining proteins bound to GSH beads after thrombin cleavage. *Purified PFK2* lanes are proteins eluted from the GSH beads with thrombin cleavage. *Purified PFK2 + ATP* are proteins incubated with ATP alone. *Purified PFK2 + ATP + PKA* are proteins phosphorylated by PKA. (**C**) Western blot analysis of GST-PFKFB2 and mutant proteins showing they are phosphorylated (Ser483) upon extraction from bacteria (*left lanes*), lose phosphorylation after purification (*middle lanes*), but are re-phosphorylated upon treatment with PKA and ATP (*right lanes*). (**D**) Representative western blot against phosphorylated-PFKFB2 (Ser483) of bacterial samples (WT PFKFB2) treated/not treated with λ-phosphatase for indicated times before lysis. Equal amounts of purified PFKFB2 were treated and loaded into each lane. Quantitation of bacterial samples treated with λ-phosphatase. Data are shown as mean±SD. The original, unedited blots are included in **Supplemental Figure 2**.

### The phosphorylation and dephosphorylation of PFKFB2 is not mediated via autonomous mechanisms

Bacteria are largely considered deficient in serine/threonine kinases, thus raising the possibility that PFKFB2 autophosphorylates. Indeed, several metabolic small-molecule kinases can “moonlight” as protein kinases ^21^. We therefore sought to determine if intrinsic kinase activity of PFKFB2 was responsible for its *in situ* phosphorylation. Previous studies identified Lys174 within the ATP binding site as essential for kinase activity ^1,22^. We therefore mutated this critical residue to glycine (K174G; **Figure 1A&C**). Interestingly, this mutation caused no change in the phosphorylation status of the protein upon purification (**Figure 1A&C)**. In addition, we created a phosphatase null PFKFB2 by mutating a critical histidine to alanine (H259A) in the phosphatase catalytic site ^11^, and a double mutant (K174G and H259A) lacking both kinase and phosphatase activities. In this manner, we could also test whether dephosphorylation of PFKFB2 was also occurring in an autonomous manner. All three mutants expressed comparably and were phosphorylated *in situ* during bacterial expression. **Figure 1C** shows the GST-fusion proteins prior to and after thrombin cleavage. All constructs dephosphorylated during purification (**Figure 1C** *middle lanes*; -PKA) and could be rephosphorylated by PKA (**Figure 1C**).

We next treated PFKFB2 (WT, K174G, H259A and double mutant) with a broad-specificity phosphatase (protein phosphatase lambda) for increasing durations to further validate the phosphorylation status (**Figure 1D**). These experiments were performed on crude bacterial lysates as this largely preserved the phosphorylation status of PFKFB2 recombinant protein. Western blot revealed PFKFB2 dephosphorylation occurred in a time-dependent manner in the presence of phosphatase (**Figure 1D**). The rate and magnitude of dephosphorylation was similar between WT and the 3 mutants (**Figure 1D**). In the absence of phosphatase, dephosphorylation was sporadic but occurred to a similar in magnitude between WT and the 3 mutants (**Figure 1D**). Cumulatively, these results support that PFKFB2 is phosphorylated in bacteria by an endogenous serine protein kinase activity ^23^.

### Identification of PFKFB2 modulators using a coupled kinase activity assay

Previous studies have used phosphofructokinase (PFK) isolated from potato tubers to measure fructose-2,6-bisphosphate (F-2,6-BP) levels ^18^. The premise is that this plant isoform of PFK (PPi-PFK) PPi-PFK1 activity is proportional to the amount of F-2,6-BP present. The assay is ultimately coupled to glycerol-3-phosphate dehydrogenase (GPDH) activity and the oxidation of NADH, monitored spectrophotometrically at 340nm (**Figure 2A**). Here, we optimized this coupled assay to use as a readout of PFKFB2 kinase activity. As validation, the kinase-null mutant of PFKFB2 (K174G) was compared to WT PFKFB2. We found that K174G kinase activity was negligible (less than 5%) as compared to WT (not shown).

**Figure 2:**
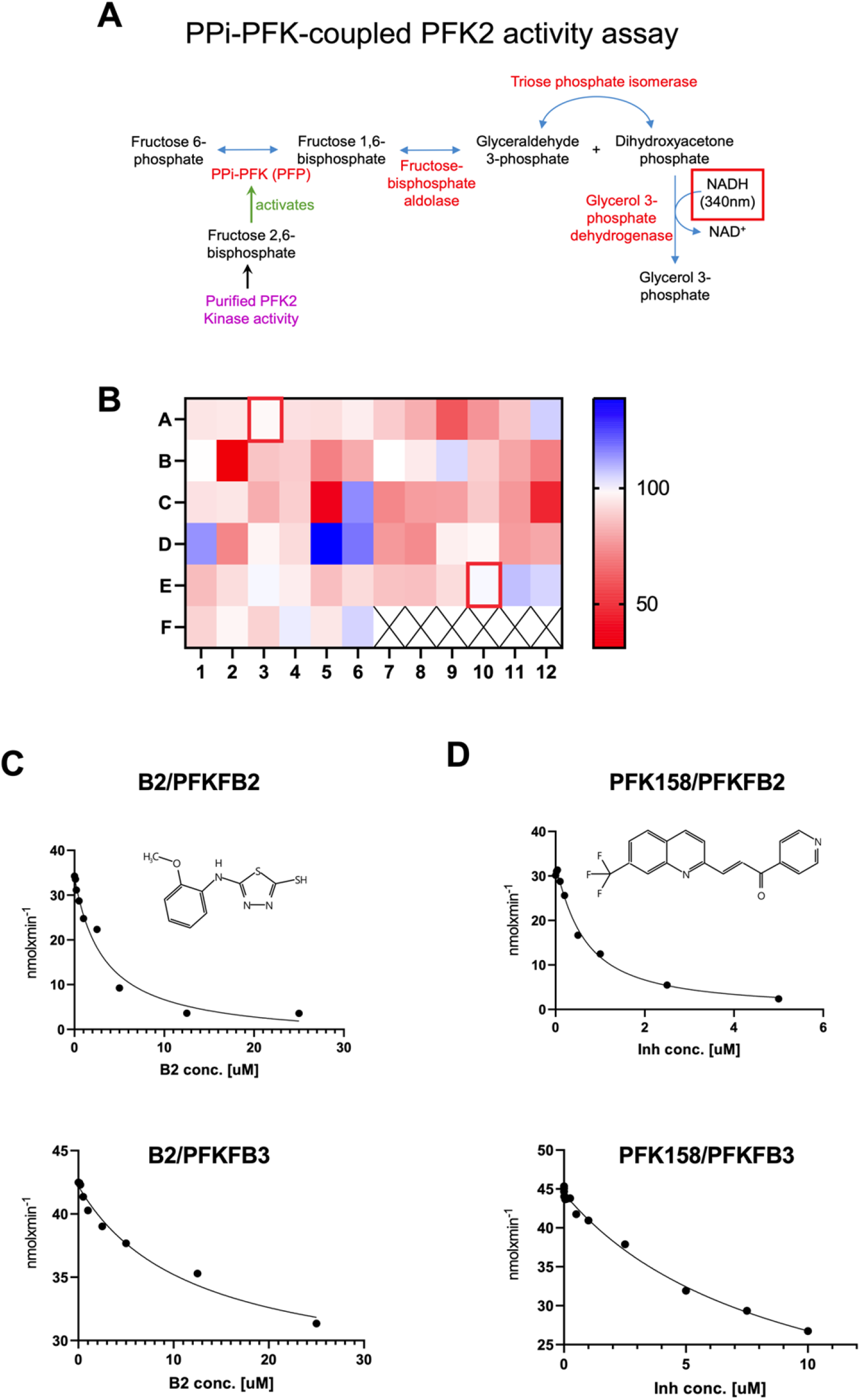
Development of a PFKFB kinase activity assay and screening of potential modulators. (**A**) Schematic of the coupled enzyme activity assay protocol. *Purple* indicates the reaction mix containing purified PFKFB isoforms. *Red* indicates enzymes added for the coupled assay. The ultimate product is glycerol-3-phosphate and the assay measures the disappearance of NADH at 340nm. (**B**) Plate-reader screening of candidate compounds (all at 10 μM in DMSO; wells A3 and E10 (*red boxes*) were DMSO controls). (**C**) IC50 determination of candidate inhibitor B2 (structure on *inset*) for PFKFB2 (*top*) and PFKFB3 (*lower*) kinase activities. (**D**) IC50 determination of known PFKFB3 kinase inhibitor PFK158 (structure on *inset*) for PFKFB2 (*top*) and PFKFB3 (*lower*) kinase activities.

We next sought to identify small molecule activators and/or inhibitors of PFKFB2. Putative molecules were selected using the AtomNet® technology (Atomwise, San Francisco California ^10^) that predicts small molecule binders using a combination of AI and high-resolution protein crystallography data (obtained from ^15^). Sixty-four compounds were tested, each at 10 μM final concentration. A heat map, generated from the rates of WT PFKFB2 activity in the presence of testing compounds, demonstrates that several compounds were able to increase (*blue*) or inhibit (*red)* PFKFB2 activity as compared to DMSO control (**Figure 2B**). The strongest activator (compound D5) was obtained from a commercial source and investigated for further validation but failed to replicate the activation seen in the initial screen. We therefore focused on further characterizing a potential new inhibitor of PFKFB2 activity.

### Validation of B2 as a new inhibitor of PFKFB2

The most potent inhibitor of PFKFB2 activity was in well B2 (**Figure 2B**). This compound was unblinded as 5-[(2-methoxyphenyl)amino]-1,3,4-thiadiazole-2-thiol but referred to for simplicity as B2 in this work. See inset on **Figure 2C** for the molecular structure. This molecule has not been previously described as an inhibitor of PFKFB2 kinase activity. A concentration course of B2 confirmed it inhibited PFKFB2 kinase activity with an IC_50_ of 3.289 μM (**Figure 2C**). We next tested whether B2 has specificity towards the PFKFB2 isoform or if it broadly inhibits PFKFB kinases. Recombinant PFKFB3 was expressed and purified (**Supplement Figure 1A**). A concentration course of B2 demonstrated it inhibited PFKFB3 as well with an IC_50_ of 11.89 μM (**Figure 2C**). Other inhibitors of PFKFB kinase activity have been previously described but they have distinctly different chemical structures from B2. We next sought to determine how B2 compares to one such established PFKFB inhibitor, PFK158 ^24,25^ (structure shown in **Figure 2D**) a derivative of the first characterized inhibitor 3PO ^26^. Under the same experimental enzyme assay conditions, a concentration course of PFK158 on recombinant PFKFB2 and PFKFB3 revealed IC_50_’s of 0.629 μM and 8.939 μM, respectively (**Figure 2D**). Thus, B2 is slightly less potent than PFK158 but these data validate B2 as an inhibitor of PFKFB2 and PFKFB3 kinase activities.

### B2 decreases cellular glycolysis

We next examined whether B2 can affect cellular glycolysis and metabolism. We chose the A-498 (HTB44) cancer cell line because of its high rate of glycolysis and its expression of both PFKFB2 and PFKFB3 isoforms (Ref. ^19^ and confirmed by western blot as shown in **Supplement Figure 1B**). Seahorse XFe24 analysis of extracellular acidification rates (ECAR) revealed that both 5.0 and 25 uM B2 decreased basal glycolysis similarly. Furthermore, the magnitude of inhibition was similar to that induced by 5.0 uM PFK158 (**Figure 3A**). However, there were differences between the inhibitors upon their effects on glycolytic capacity (**Figure 3B**). Glycolytic capacity is the maximal rate of glycolysis achieved upon the addition of oligomycin. Both 5.0 and 25 uM B2, but not PFK158, inhibited glycolytic capacity significantly (**Figure 3B**).

**Figure 3:**
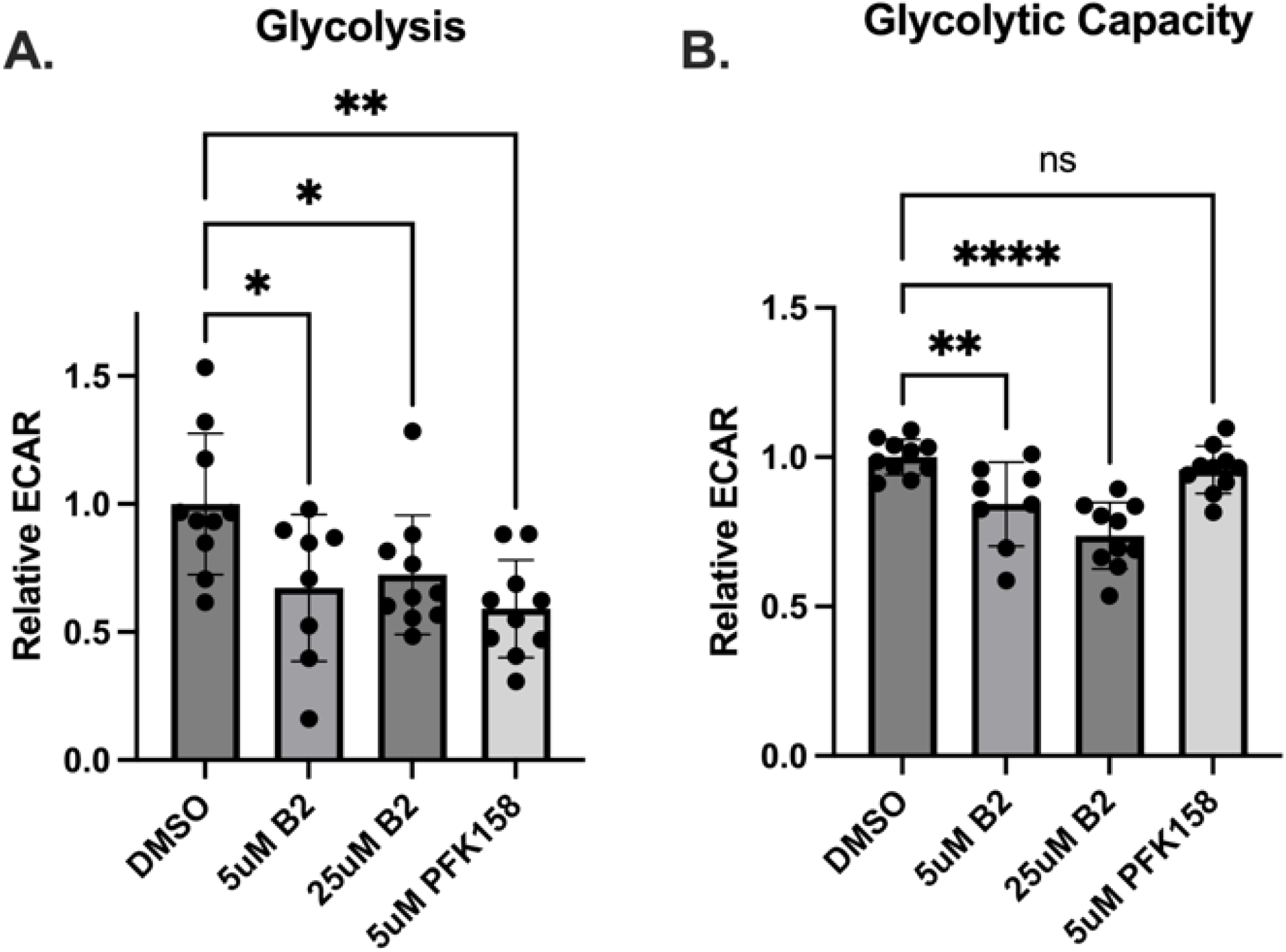
B2 inhibits glycolysis and glycolytic capacity. A-498 cells were treated with 5μM B2 for 60 minutes. The rates of glycolysis (**A**) were measured as ECAR upon addition of glucose and glycolytic capacity (**B**) as the rate of ECAR after the subsequent addition of oligomycin. Each point represents an individual well from a Seahorse XFe24 plate, with the experiment performed on 2 separate plates. Data are shown as mean±SD. Not significant (ns): *P*>0.05, \**P*≤0.05, \*\**P*≤0.01, \*\*\*\**P*≤0.0001by one-way ANOVA with multiple comparison of the mean of each test group to the mean of the DMSO control.

We next performed LC-MS targeted metabolomic analysis on intermediates extracted from A-498 cells that were treated with either 5.0 uM B2 or PFK158 for 60 min. The targeted analysis focused on glycolytic intermediates, but included other select Krebs cycle and amino acids (14 metabolites were quantified). F-2,6-BP itself was not measured, as it is challenging to detect due to its low abundance, instability, and the lack of commercially available standards. As shown in **Figure 4A**, the differential effects of B2 and PFK158 on cellular metabolism are seen in principal component analysis (**Figure 4**). All groups were distinct, demonstrating unique, nonoverlapping effects of B2 and PFK158 on cellular metabolism. Examining specific glycolytic intermediates, PFK158, but not B2, caused a significant decrease in glucose-6-phosphate.

**Figure 4:**
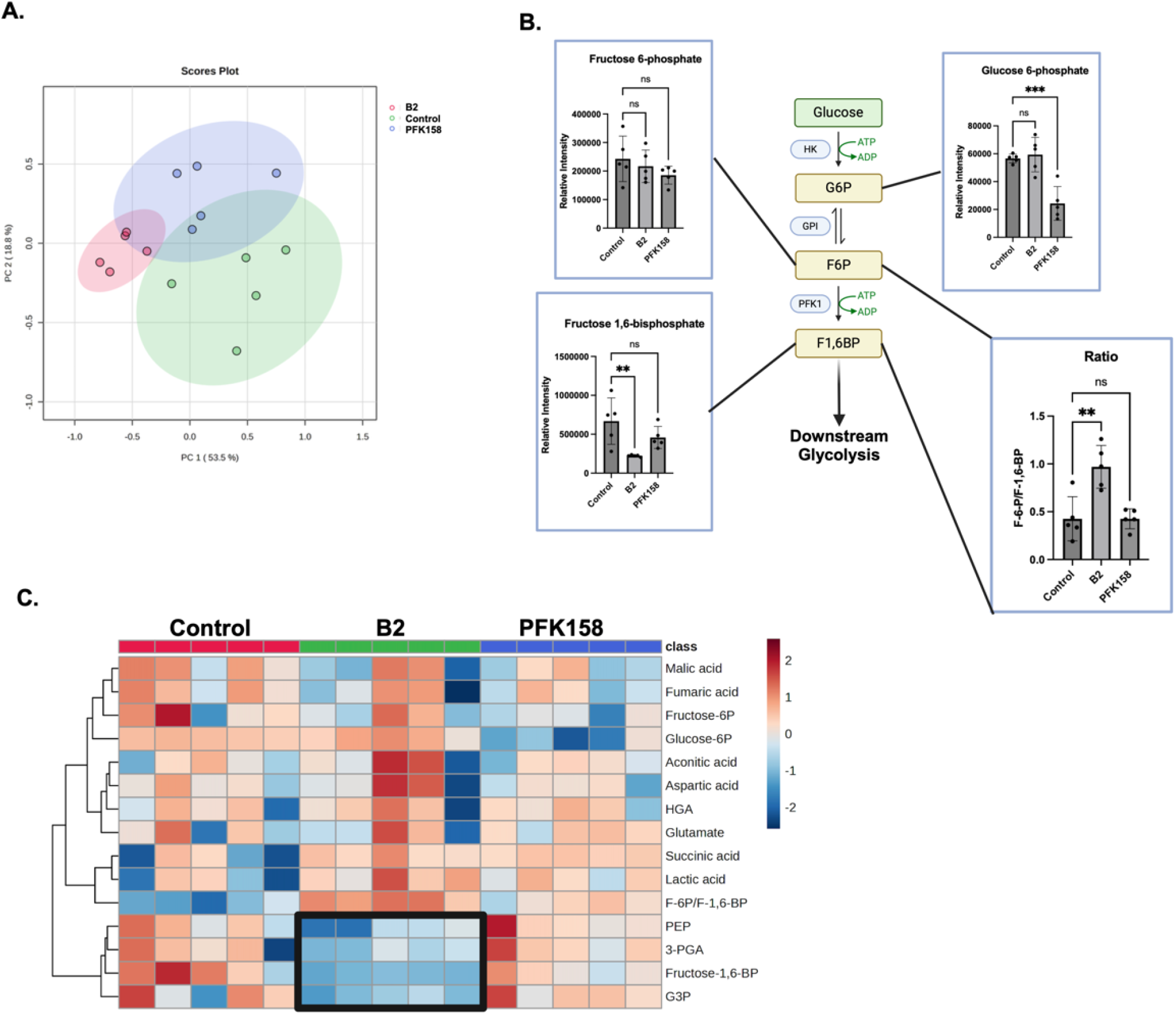
B2 decreases glycolytic intermediates in cells. A-498 cells were treated with B2 or PFK158 for 60 min and then metabolites were extracted for LC/MS analysis. (**A**) PCA analysis of control, B2 and PFK158 treatment groups. (**B**) Early glycolytic intermediates and the ratio of F-6-P to F-1,6-BP (*lower right*) are shown. **(C)** Heatmap of targeted metabolomic data. Each column is a separate biological replicate (n=5). The box indicates the clustering of decreased downstream glycolytic intermediates in B2-treated cells. The ratio of fructose-6-phosphate (F-6P) to fructose-1,6-bisphosphate (F-1,6-BP) is included in the heatmap. Other abbreviations: HGA, DL-hydroxyglutaric acid; PEP, phosphoenolpyruvic acid; 3-PGA, 3-phosphoglyceric acid; G3P; glyceraldehyde 3-phosphate; Data (n=5) are shown as mean±SD. Not significant (ns): \*\**P*≤0.01, \*\*\*\**P*≤0.0001by one-way ANOVA with multiple comparison of the mean of each test group to the mean of the vehicle control. PCA and heatmap were generated with MetaboAnalyst 6.0; figure produced with Biorender.

Neither compound affected levels of fructose-6-phosphate. However, only B2 caused a significant decrease in fructose-1,6-bisphosphate and the ratio of fructose-6-phosphate to fructose-1,6-bisphosphate (**Figure 4B**). This supports that B2 is inhibiting PFKFBs activation of glycolysis at the PFK1 step of glycolysis. A heatmap of metabolites show that B2 treatment, but not PFK158, had decreases in glycolytic intermediates downstream of PFK-1 enzyme activity relative to control cells (**Figure 4C**, *boxed area*). These intermediates included phosphoenolpyruvate (PEP), 3-phosphoglyceric acid (3-PGA), and glyceraldehyde 3-phosphate (G3P). In conclusion, these results demonstrate B2 is a glycolytic inhibitor with cellular metabolic bioactivity that is distinct from known PFKFB inhibitor, PFK158.

## DISCUSSION

The goals of the present study were multifold. First, we sought to express recombinant PFKFB2 and develop a robust protocol to measure its kinase activity. Second, upon successful completion, we tested a library of small molecules, selected by a robust AI-platform, that target PFKFB2 kinase activity. Third, the study resulted in the identification of a new inhibitor that was further validated on both PFKFB2 and PFKFB3 kinase activities. Finally, we showed that this newly identified PFKFB2 inhibitor decreases cellular glycolytic rates.

An unanticipated result of the study was the finding that PFKFB2 expressed in bacteria is phosphorylated on Ser483. This is a well-characterized phosphorylation site that increases PFKFB2 kinase activity at the expense of decreased phosphatase activity. Bacteria are largely considered devoid of serine/threonine kinase activity, thus raising the intriguing possibility that PFKFB2 can autophosphorylate by “moonlighting” as a protein kinase. This line of reasoning is supported by work showing that PFKFB4 can regulate transcriptional reprogramming by phosphorylating and activating the oncogenic steroid receptor coactivator-3 (SRC-3) protein ^27^. However, our investigations revealed this is not likely the case with PFKFB2. A kinase-null mutant was generated and purified from bacteria still emerged phosphorylated at Ser483. This supports the more recent conclusions that bacteria do indeed contain serine/threonine kinase activities ^23^. Furthermore, the dephosphorylation of PFKFB2 is labile but not likely due to intrinsic protein phosphatase activity. A phosphatase-null mutant of PFKFB2 still dephosphorylated in a time dependent manner upon purification.

The measurement of PFKFB2 kinase activity is challenging. The product, F-2,6-BP, is not stable and its in vivo concentration is low. Mass spectrometry detection of F-2,6-BP is further complicated by the lack of a commercially available standard. The current protocol for measuring F-2,6-BP is through a coupled assay that relies on the activation of PPi-PFK purified from potato tubers ^18^. We have previously adapted this assay for measurement of F-2,6-BP levels in mouse hearts ^17^ and here we have refined the assay to measure the activity of recombinant PFKFB2 kinase activity. Validation of the assay was achieved by showing that the kinase-null mutant of PFKFB2 lacked detectable activity. This allowed us to then screen small molecules that could potentially modulate PFKFB2 kinase activity.

Our initial goal was to seek out potential small molecules that could enhance PFKFB2 kinase activity because we have previously shown that PFKFB2 expression and activation are decreased in the diabetic heart ^28,29^ and increasing its kinase activity has protective effects against high fat diet-induced dysfunction ^30,31^. To do so, Atomwise used their AtomNet® technology to target the bisphosphatase site, testing the hypothesis that inhibiting phosphatase activity may reciprocally enhance kinase activity of the bifunctional enzyme. For the virtual screen, the X-ray crystal structure of the human PFKFB2 dimer was used (PDB ID: 5HTK; ^15^). PFKFB2 functions as a dimer where each bifunctional monomer is composed of four regions, with the core kinase and bisphosphatase domains at the center and 2 regulatory regions at the termini. The terminal regions regulate the enzyme’s function by changing tertiary or quaternary conformations in response to several effectors ^3^ but are notably missing from the crystal structure. Sequence alignment of the human and mouse PFKFB2 shows a near conservation (∼90% identity) of residues forming the bisphosphatase domain (amino acids 249 – 505).

Therefore, identified modulators have a high probability of targeting both human and mouse PFKFB2. Our initial screen of small molecules showed three potential activators but failed further validation. Nevertheless, our screen also showed the presence of several potential inhibitors. Despite targeting the bisphosphatase site, the identification of potential kinase inhibitors was not completely unanticipated. The binding of small molecules to discrete sites distal to the kinase domain may have unanticipated effects on the structure and enzymatic activities. Furthermore, the identification of new PFKFB kinase inhibitors was welcome because of their potential clinical importance. The strongest inhibitor, B2, was therefore subjected to further characterize.

A potential limitation of our PFKFB kinase activity screen is that it is based upon a coupled assay. It is therefore possible that the small molecules being screened are interfering with one or more of the coupled enzymes (see **Figure 2A**). To test this possibility, we extracted F-2,6-BP from the hearts of Glyco^Hi^ mice. This is a transgenic mouse model that overexpresses a mutated form of PFKFB1 in the heart that has enhanced kinase activity and subsequently increased cardiac glycolysis ^32^. We have previously used the coupled assay employed here to show that Glyco^Hi^ mouse hearts have persistently elevated F-2,6-BP levels ^17^. In control experiments, we found that B2 had negligible effects on F-2,6-BP measurements from Glyco^Hi^ mouse hearts (not shown). This supports the conclusion that B2 is directly affecting PFKFB kinase activity.

The PFKFB family of enzymes, and especially PFKFB3, are sought after targets for inhibition of kinase activity. This is because of their potential to specifically target cancer types that upregulate expression of PFKFB3 to increase glycolysis. The first and most widely used inhibitor is 3-(3-pyridinyl)-1-(4-pyridinyl)-2-propen-1-one (3PO) ^26^, while more recently other inhibitors based upon 3PO have been developed including PFK158 ^4^. As shown in **Figures 2C** and **2D**, B2 and PFK158 are chemically distinct. Of note, B2 has a free thiol group, which may be important for its inhibitory activity. Compounds with free thiol groups can dimerize under non-physiological high pH when cyclic azo groups deprotonate. In addition, thiol groups can react with cysteine residues on proteins to form covalent adducts. However, such reactions are relatively slow and in our experiments no time dependence was observed for the inhibition of PFKFB2. Furthermore, the addition of a thiol-reducing agent had no effect on inhibition and supports covalent binding of B2 to the protein is unlikely.

With the growing identification of the importance of PFKFB isoforms in health and disease, modulators of their kinase activity may have broad biological importance. This includes the potential for applications outside of cancer, including myocardial ischemia ^33^, sepsis ^6^, fibrosis ^7^, and obesity ^8^. One caveat of the current study is that we cannot eliminate the possibility that B2 is direct affecting PFK1 or other glycolytic enzymes in our cell-based assays. Future studies will further investigate the effects of B2 on cellular metabolism and determine its potential application in targeting cells with high glycolytic capacity. Furthermore, this study opens the possibility for further investigation and refinement to more selectively target specific PFKFB isoforms.

## ACKNOWLEDGEMENTS

This work was supported by the Atomwise AIMS program (Award A18-142) and the National Institutes of Health grant R01HL160955 (Humphries). Metabolomic data was generated by the Metabolic Phenotyping Core, supported by P20GM139763.

## DATA AVAILABILITY

The corresponding author (K.M.H.) can be contacted for any data availability requests.

## REFERENCES

1 Rider, M. H. et al. 6-phosphofructo-2-kinase/fructose-2,6-bisphosphatase: head-to-head with a bifunctional enzyme that controls glycolysis. Biochem J 381, 561–579 (2004). 10.1042/BJ20040752

2 Bartrons, R. et al. Fructose 2,6-Bisphosphate in Cancer Cell Metabolism. Front Oncol 8, 331 (2018). 10.3389/fonc.2018.00331

3 Okar, D. A. et al. PFK-2/FBPase-2: maker and breaker of the essential biofactor fructose-2,6-bisphosphate. Trends Biochem Sci 26, 30–35 (2001). 10.1016/s0968-0004(00)01699-6

4 Jones, B. C., Pohlmann, P. R., Clarke, R. & Sengupta, S. Treatment against glucose-dependent cancers through metabolic PFKFB3 targeting of glycolytic flux. Cancer Metastasis Rev 41, 447–458 (2022). 10.1007/s10555-022-10027-5

5 Kitamura, K., Kangawa, K., Matsuo, H. & Uyeda, K. Phosphorylation of myocardial fructose-6-phosphate,2-kinase: fructose-2,6-bisphosphatase by cAMP-dependent protein kinase and protein kinase C. Activation by phosphorylation and amino acid sequences of the phosphorylation sites. J Biol Chem 263, 16796–16801 (1988).

6 Xiao, M., Liu, D., Xu, Y., Mao, W. & Li, W. Role of PFKFB3-driven glycolysis in sepsis. Ann Med 55, 1278–1289 (2023). 10.1080/07853890.2023.2191217

7 Liu, Q. et al. Advances in the understanding of the role and mechanism of action of PFKFB3-mediated glycolysis in liver fibrosis (Review). Int J Mol Med 54 (2024). 10.3892/ijmm.2024.5429

8 Griesel, B. A. et al. PFKFB3-dependent glucose metabolism regulates 3T3-L1 adipocyte development. FASEB J 35, e21728 (2021). 10.1096/fj.202100381RR

9 Yi, M. et al. 6-Phosphofructo-2-kinase/fructose-2,6-biphosphatase 3 and 4: A pair of valves for fine-tuning of glucose metabolism in human cancer. Mol Metab 20, 1–13 (2019). 10.1016/j.molmet.2018.11.013

10 Atomwise, A. P. AI is a viable alternative to high throughput screening: a 318-target study. Sci Rep 14, 7526 (2024). 10.1038/s41598-024-54655-z

11 Argaud, D. et al. Adenovirus-mediated overexpression of liver 6-phosphofructo-2-kinase/fructose-2,6-bisphosphatase in gluconeogenic rat hepatoma cells. Paradoxical effect on Fru-2,6-P2 levels. J Biol Chem 270, 24229–24236 (1995). 10.1074/jbc.270.41.24229

12 Wallach, I., Dzamba, M. & Heifets, A. AtomNet: a deep convolutional neural network for bioactivity prediction in structure-based drug discovery. arXiv preprint arXiv:1510.02855 (2015).

13 Hsieh, C. H. et al. Miro1 Marks Parkinson’s Disease Subset and Miro1 Reducer Rescues Neuron Loss in Parkinson’s Models. Cell Metab 30, 1131–1140e1137 (2019). 10.1016/j.cmet.2019.08.023

14 Su, S. et al. SPOP and OTUD7A Control EWS-FLI1 Protein Stability to Govern Ewing Sarcoma Growth. Adv Sci (Weinh) 8, e2004846 (2021). 10.1002/advs.202004846

15 Crochet, R. B. et al. Crystal structure of heart 6-phosphofructo-2-kinase/fructose-2,6-bisphosphatase (PFKFB2) and the inhibitory influence of citrate on substrate binding. Proteins 85, 117–124 (2017). 10.1002/prot.25204

16 Bruns, R. F. & Watson, I. A. Rules for identifying potentially reactive or promiscuous compounds. J Med Chem 55, 9763–9772 (2012). 10.1021/jm301008n

17 Batushansky, A. et al. GC-MS metabolic profiling reveals fructose-2,6-bisphosphate regulates branched chain amino acid metabolism in the heart during fasting. Metabolomics 15, 18 (2019). 10.1007/s11306-019-1478-5

18 Van Schaftingen, E., Lederer, B., Bartrons, R. & Hers, H. G. A kinetic study of pyrophosphate: fructose-6-phosphate phosphotransferase from potato tubers. Application to a microassay of fructose 2,6-bisphosphate. Eur J Biochem 129, 191–195 (1982). 10.1111/j.1432-1033.1982.tb07039.x

19 Li, J. et al. Overexpression of PFKFB3 promotes cell glycolysis and proliferation in renal cell carcinoma. BMC Cancer 22, 83 (2022). 10.1186/s12885-022-09183-2

20 Rider, M. H. et al. Evidence for new phosphorylation sites for protein kinase C and cyclic AMP-dependent protein kinase in bovine heart 6-phosphofructo-2-kinase/fructose-2,6-bisphosphatase. FEBS Lett 310, 139–142 (1992). 10.1016/0014-5793(92)81315-d

21 Lu, Z. & Hunter, T. Metabolic Kinases Moonlighting as Protein Kinases. Trends Biochem Sci 43, 301–310 (2018). 10.1016/j.tibs.2018.01.006

22 Bertrand, L. et al. Site-directed mutagenesis of Lys-174, Asp-179 and Asp-191 in the 2-kinase domain of 6-phosphofructo-2-kinase/fructose-2,6-bisphosphatase. Biochem J 321 (Pt 3), 623–627 (1997). 10.1042/bj3210623

23 Pereira, S. F., Goss, L. & Dworkin, J. Eukaryote-like serine/threonine kinases and phosphatases in bacteria. Microbiol Mol Biol Rev 75, 192–212 (2011). 10.1128/MMBR.00042-10

24 Mondal, S. et al. Therapeutic targeting of PFKFB3 with a novel glycolytic inhibitor PFK158 promotes lipophagy and chemosensitivity in gynecologic cancers. Int J Cancer 144, 178–189 (2019). 10.1002/ijc.31868

25 Clem, B. F. et al. Targeting 6-phosphofructo-2-kinase (PFKFB3) as a therapeutic strategy against cancer. Mol Cancer Ther 12, 1461–1470 (2013). 10.1158/1535-7163.MCT-13-0097

26 Clem, B. et al. Small-molecule inhibition of 6-phosphofructo-2-kinase activity suppresses glycolytic flux and tumor growth. Mol Cancer Ther 7, 110–120 (2008). 10.1158/1535-7163.MCT-07-0482

27 Dasgupta, S. et al. Metabolic enzyme PFKFB4 activates transcriptional coactivator SRC-3 to drive breast cancer. Nature 556, 249–254 (2018). 10.1038/s41586-018-0018-1

28 Bockus, L. B. & Humphries, K. M. cAMP-dependent Protein Kinase (PKA) Signaling Is Impaired in the Diabetic Heart. J Biol Chem 290, 29250–29258 (2015). 10.1074/jbc.M115.681767

29 Bockus, L. B. et al. Cardiac Insulin Signaling Regulates Glycolysis Through Phosphofructokinase 2 Content and Activity. J Am Heart Assoc 6 (2017). 10.1161/JAHA.117.007159

30 Newhardt, M. F. et al. Enhancing cardiac glycolysis causes an increase in PDK4 content in response to short-term high-fat diet. J Biol Chem 294, 16831–16845 (2019). 10.1074/jbc.RA119.010371

31 Mendez Garcia, M. F. et al. Increased cardiac PFK-2 protects against high-fat diet-induced cardiomyopathy and mediates beneficial systemic metabolic effects. iScience 26, 107131 (2023). 10.1016/j.isci.2023.107131

32 Wang, Q. et al. Cardiac phosphatase-deficient 6-phosphofructo-2-kinase/fructose-2,6-bisphosphatase increases glycolysis, hypertrophy, and myocyte resistance to hypoxia. Am J Physiol Heart Circ Physiol 294, H2889–2897 (2008). 10.1152/ajpheart.91501.2007

33 Yang, Q. et al. PFKFB3 Inhibitor 3PO Reduces Cardiac Remodeling after Myocardial Infarction by Regulating the TGF-beta1/SMAD2/3 Pathway. Biomolecules 13 (2023). 10.3390/biom13071072

